# Fast and Stable Signal Deconvolution via Compressible State-Space Models

**DOI:** 10.1101/092643

**Authors:** Abbas Kazemipour, Ji Liu, Krystyna Solarana, Daniel A. Nagode, Patrick O. Kanold, Min Wu, Behtash Babadi

**Author notes:** Corresponding authors: A. Kazemipour and Behtash Babadi. This work has been presented in part at the 2016 IEEE Global Conference on Signal and Information Processing as an invited paper [1].

## Abstract

Common biological measurements are in the form of noisy convolutions of signals of interest with possibly unknown and transient blurring kernels. Examples include EEG and calcium imaging data. Thus, signal deconvolution of these measurements is crucial in understanding the underlying biological processes. The objective of this paper is to develop fast and stable solutions for signal deconvolution from noisy, blurred and undersampled data, where the signals are in the form of discrete events distributed in time and space. *Methods*: We introduce compressible state-space models as a framework to model and estimate such discrete events. These state-space models admit abrupt changes in the states and have a convergent transition matrix, and are coupled with compressive linear measurements. We consider a dynamic compressive sensing optimization problem and develop a fast solution, using two nested Expectation Maximization algorithms, to jointly estimate the states as well as their transition matrices. Under suitable sparsity assumptions on the dynamics, we prove optimal stability guarantees for the recovery of the states and present a method for the identification of the underlying discrete events with precise confidence bounds. *Results*: We present simulation studies as well as application to calcium deconvolution and sleep spindle detection, which verify our theoretical results and show significant improvement over existing techniques. *Conclusion*: Our results show that by explicitly modeling the dynamics of the underlying signals, it is possible to construct signal deconvolution solutions that are scalable, statistically robust, and achieve high temporal resolution. *Significance*: Our proposed methodology provides a framework for modeling and deconvolution of noisy, blurred, and undersampled measurements in a fast and stable fashion, with potential application to a wide range of biological data.

## I. Introduction

In many signal processing applications such as estimation of brain activity from magnetoencephalography (MEG) time-series [2], estimation of time-varying networks [3], electroencephalogram (EEG) analysis [4], calcium imaging [5], functional magnetic resonance imaging (fMRI) [6], and video compression [7], the signals often exhibit abrupt changes that are blurred through convolution with unknown kernels due to intrinsic measurement constraints. Extracting the underlying signals from blurred and noisy measurements is often referred to as signal deconvolution. Traditionally, state-space models have been used for such signal deconvolution problems, where the states correspond to the unobservable signals. Gaussian state-space models in particular are widely used to model smooth state transitions. Under normality assumptions, posterior mean filters and smoothers are optimal estimators, where the analytical solution is given respectively by the Kalman filter and the fixed interval smoother [8], [9].

When applied to observations from abruptly changing states, Gaussian state-space models exhibit poor performance in recovering sharp transitions of the states due to their underlying smoothing property. Although filtering and smoothing recursions can be obtained in principle for non-Gaussian state-space models, exact calculations are no longer possible. Apart from crude approximations like the extended Kalman filter, several methods have been proposed including numerical methods for low-dimensional states [10], Monte Carlo filters [10], [11], posterior mode estimation [12], [13], and fully Bayesian smoothing using Markov chain Monte Carlo simulation [14], [15]. In order to exploit sparsity, several dynamic compressed sensing (CS) techniques, such as the Kalman filtered CS algorithm, have been proposed that typically assume partial information about the sparse support or estimate it in a greedy and online fashion [16], [17], [18], [19], [20]. However, little is known about the theoretical performance guarantees of these algorithms.

In this paper, we consider the problem of estimating state dynamics from noisy and undersampled observations, where the state transitions are governed by autoregressive models with compressible innovations. Motivated by the theory of CS, we employ an objective function formed by the *ℓ*_1_-norm of the state innovations [21]. Unlike the traditional compressed sensing setting, the sparsity is associated with the dynamics and not the states themselves. In the absence of observation noise, the CS recovery guarantees are shown to extend to this problem [21]. However, in a realistic setting in the presence of observation noise, it is unclear how the CS recovery guarantees generalize to this estimation problem.

We will present stability guarantees for this estimator under a convergent state transition matrix, which confirm that the CS recovery guarantees can be extended to this problem. The corresponding optimization problem in its Lagrangian form is akin to the MAP estimator of the states in a linear state-space model where the innovations are Laplace distributed. This allows us to integrate methods from Expectation-Maximization (EM) theory and Gaussian state-space estimation to derive efficient algorithms for the estimation of states as well as the state transition matrix, which is usually unknown in practice. To this end, we construct two nested EM algorithms in order to jointly estimate the states and the transition matrix. The outer EM algorithm for state estimation is akin to the fixed interval smoother, and the inner EM algorithm uses the state estimates to update the state transition matrix [22]. The resulting EM algorithm is recursive in time, which makes the computational complexity of our method scale linearly with temporal dimension of the problem. This provides an advantage over existing methods based on convex optimization, which typically scale super-linearly with the temporal dimension.

Our results are related to parallel applications in spectral estimation, source localization, and channel equalization [23], where the measurements are of the form **Y** = **AX** + **N**, with **Y** is the observation matrix, **X** denotes the unknown parameters, **A** is the measurement matrix, and **N** is the additive noise. These problems are referred to as Multiple Measurement Vectors (MMV) [24] and Multivariate Regression [25]. In these applications, solutions with row sparsity in **X** are desired. Recovery of sparse signals with Gaussian innovations is studied in [26]. Several recovery algorithms including the *ℓ*_1_–ℓ_q_ minimization methods, subspace methods and greedy pursuit algorithms [27] have been proposed for support union recovery in this setup. Our contributions are distinct in that we directly model the state innovations as a compressible sequence, for recovery of which we present both sharp theoretical guarantees as well as fast algorithms from state-space estimation.

Finally, we provide simulation results as well as applications to two experimentally-acquired data sets: calcium imaging recordings of neuronal activity, and EEG data during sleep. In the former, the deconvolution problem concerns estimating the location of spikes given the temporally blurred calcium fluorescence, and in the latter, the objective is to detect the occurrence and onset of sleep spindles. Our simulation studies confirm our theoretical predictions on the performance gain obtained by compressible state-space estimation over those obtained by traditional estimators such as the basis pursuit denoising. Our real data analyses reveal that our compressible state-space modeling and estimation framework outperforms two of the commonly-used methods for calcium deconvolution and sleep spindle detection. In the spirit of easing reproducibility, we have made the MATLAB implementation of our algorithm publicly available [28].

The rest of this paper is organized as follows. In the following section, we formulate compressible state-space models, introduce our notation and present our main theoretical analysis on the stability of the state estimation from undersampled measurements. In Section II-B, we introduce a fast solver for joint estimation of states and their transition matrices, which we have named FCSS. We provide simulation studies and application to calcium deconvolution and sleep spindle detection in Section III. We discuss the implication of our results in Section IV, followed by concluding remarks in Section V.

## II. Methods

In this section, we establish our problem formulation and notational conventions and present our main theoretical analysis and algorithm development. In the interest of space, the description of the experimental procedures is given in the supplementary material.

### A. Problem Formulation and Theoretical Analysis

Throughout the paper we use bold lower and upper case letters for denoting vectors and matrices, respectively. We denote the support of a vector x_*t*_∈ℝ^*p*^ by supp(x_*t*_) and its *j*th element by (x_*t*_)_*j*_. We consider the linear compressible state-space model given by

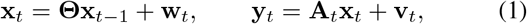

where 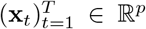 denote the sequence of unobservable states, **Θ** is the state transition matrix satisfying ∥Θ∥< 1, **w**_*t*_∈ℝ^*p*^ is the state innovation sequence, 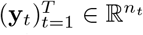 are the linear observations, **A**_*t*_∈ℝ^*n_t_*×*p*^ denotes the measurement matrix, and **e**_*t*_ ∈ℝ^*n_t_*^ denotes the measurement noise. The main problem is to estimate the unobserved sequence 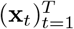 (and possibly **Θ**), given the sequence of observations 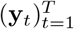 This problem is in general ill-posed, when *n_t_* < *p*, for some *t*. We therefore need to make additional assumptions in order to seek a stable solution.

Given a sparsity level *s* and a vector x, we denote the set of its *s* largest magnitude entries by *S*, and its best *s*-term approximation error by *σ*_*s*_(x) = ∥x − x_*S*_∥_1_. When *σ*_*s*_(x) ~𝒪^(1/2−**ξ**)^ for some *ξ* ≥ 0, we refer to x as (*s*, *ξ*)– compressible. We assume that the state innovations are sparse (resp. compressible), i.e. x_*t*_ − **Θ**x_*t*−1_ is *s_t_*-sparse (resp. (*s_t_*,*ξ*)- compressible) with *s*_1_ ≫ *s_t_* for *t* ∈[*T*]\{1}. Our theoretical analysis pertain to the compressed sensing regime where 1 ≪ *s_t_* < *n_t_* ≪ *p*.

For simplicity of notation, we define x_0_ to be the all-zero vector in ℝ^*p*^. For a matrix **A**, we denote restriction of **A** to its first *n* rows by (**A**)_*n*_. We say that the matrix **A** ∈ℝ^*n×p*^ satisfies the restricted isometry property (RIP) [29] of order *s*, if for all *s*-sparse x ∈ℝ^*p*^, we have

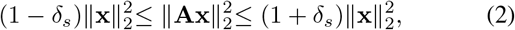

where *δ*_*s*_ ∈ (0,1) is the smallest constant for which Eq. (2) holds [30]. We assume that the rows of **A**_*t*_ are a subset of the rows of **A**_1_, i.e. **A**_*t*_ = (**A**_1_)_*n_t_*_, and define 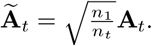 Other than its technical usefulness, the latter assumption helps avoid prohibitive storage of all the measurement matrices. In order to promote sparsity of the state innovations, we consider the dynamic *ℓ*_1_-regularization (dynamic CS from now on) problem defined as

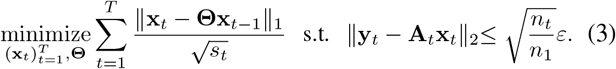

where *ε* is an upper bound on the observation noise, i.e., ∥𝑣_*t*_∥_2_ ≤ *ε* for all *t*. Note that this problem is a variant of the dynamic CS problem introduced in [21]. We also consider the modified Lagrangian form of (3) given by

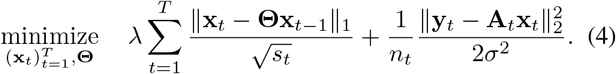

for some constants 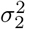 and λ ≥ 0. Note that if **v**_*t*_ ~ 𝒩(0, *n_t_𝜎*^2^**I**), then Eq. (4) is akin to the maximum *a posteriori* (MAP) estimator of the states in (1), assuming that the *state* innovations were independent Laplace random variables with respective parameters 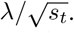 We will later use this analogy to derive fast solutions to the optimization problem in (4).

Uniqueness and exact recovery of the sequence 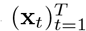 in the absence of noise was proved in [21] for **Θ** = **I**, by an inductive construction of dual certificates. The special case **Θ** = **I** can be considered as a generalization of the total variation (TV) minimization problem [31]. Our main result on stability of the solution of (3) is the following:

#### Theorem 1

(Stable Recovery of Activity in the Presence of Noise). *Let* 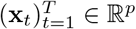 *be a sequence of states with a known transition matrix* **Θ** =𝜃**I**, *where*|𝜃|< 1 *and* Ã_*t*_,*t* ≥ 1 *satisfies RIP of order* 4*s with* 𝛿_4*s*_< 1/3. *Suppose that *n*_1_ > *n*_2_ = *n*_3_ =…= *n_T_*. *Then, the solution* 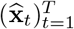 to the dynamic CS problem (3) satisfies*.

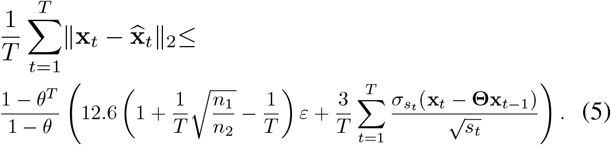

***Proof Sketch.*** The proof of Theorem 1 is based on establishing a modified cone and tube constraint for the dynamic CS problem (3) and using the boundedness of the Frobenius norm of the inverse first-order differencing operator. Details of the proof are given in Appendix **VII-A**.

#### **Remark**.

The first term on the right hand side of Theorem 1 implies that the average reconstruction error of the sequence 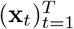 is upper bounded proportional to the noise level *ε*, which implies the stability of the estimate. The second term is a measure of compressibility of the innovation sequence and vanishes when the sparsity condition is exactly met.

### B. Fast Iterative Solution via the EM Algorithm

Due to the high dimensional nature of the state estimation problem, algorithms with polynomial complexity exhibit poor scalability. Moreover, when the state transition matrix is not known, the dynamic CS optimization problem (4) is not convex in (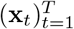,**Θ**). Therefore standard convex optimization solvers cannot be directly applied. This problem can be addressed by employing the Expectation-Maximization algorithm [22]. A related existing result considers weighted 𝓁_1_-regularization to adaptively capture the state dynamics [32]. Our approach is distinct in that we derive a fast solution to (4) via two nested EM algorithms, in order to jointly estimate the states and their transition matrix. The outer EM algorithm converts the estimation problem to a form suitable for the usage of the traditional Fixed Interval Smoothing (FIS) by invoking the EM interpretation of the Iterative Re-weighted Least Squares (IRLS) algorithms [33]. The inner EM algorithm performs state and parameter estimation efficiently using the FIS. We refer to our estimated as the Fast Compressible State-Space (FCSS) estimator.

*The outer EM loop of* FCSS: In [33], the authors established the equivalence of the IRLS algorithm as an instance of the EM algorithm for solving 𝓁_1_-minimization problems via the Normal/Independent (N/I) characterization of the Laplace distribution. Consider the 𝜖-perturbed 𝓁_1_-norm as

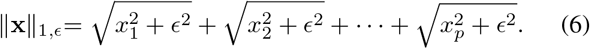

Note that for 𝜖 = 0, ∥x∥_1,𝜖_ coincides with the usual 𝓁_1_-norm. We define the 𝜖-perturbed version of the dual problem (4) by

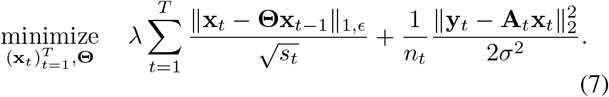

If instead of the 𝓁_1,𝜖_-norm, we had the square 𝓁_2_ norm, then the above problem could be efficiently solved using the FIS. The outer EM algorithm indeed transforms the problem of Eq. (7) into this form. Note that the 𝜖-perturbation only adds a term of the order *𝒪*(*𝜖 p*) to the estimation error bound of Theorem 1, which is negligible for small enough 𝜖 [33].

The problem of Eq. (7) can be interpreted as a MAP problem: the first term corresponds to the state-space prior 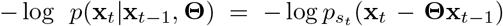, where 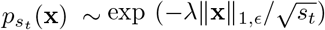 denoting the 𝜖-perturbed Laplace distribution; the second term is the negative log-likelihood of the data given the state, assuming a zero-mean Gaussian observation noise with covariance *σ*^2^**I**. Suppose that at the end of the 𝑙^th^ iteration, the estimates 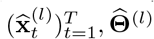 are obtained, given the observations 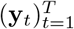. As it is shown in Appendix VII-B, the outer EM algorithm transforms the optimization problem to:

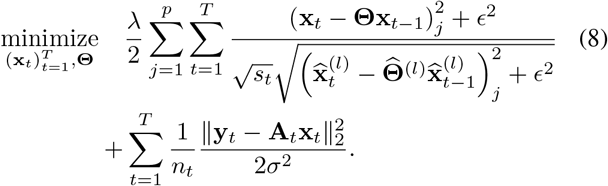

in order to find 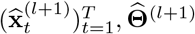. Under mild conditions, convergence of the solution of (8) to that of (4) was established in [33]. The objective function of (8) is still not jointly convex in 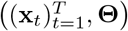. Therefore, to carry out the optimization, i.e. the outer M step, we will employ another instance of the EM algorithm, which we call the inner EM algorithm, to alternate between estimating of 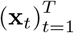 and **Θ**.

*The inner EM loop of* FCSS: Let 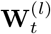 be a diagonal matrix such that

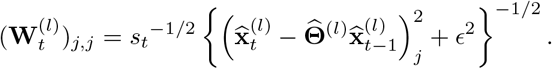

Consider an estimate 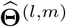, Corresponding to the *m*^th^ iteration of the inner EM algorithm within the *l*^th^ M-step of the outer EM. In this case, Eq. (8) can be thought of the MAP estimate of the Gaussian state-space model given by:

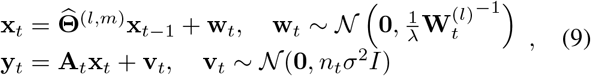

In order to obtain the inner E step, one needs to find the density of 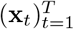 given 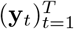 and 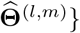. Given the Gaussian nature of the state-space in Eq. (9), this density is a multivariate Gaussian density, whose means and covariances can be efficiently computed using the FIS. For all *t* ∈ [*T*], the FIS performs a forward Kalman filter and a backward smoother to generate [34], [8]:

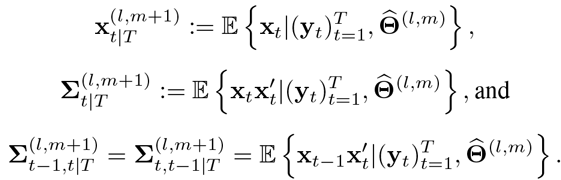

Note that due to the quadratic nature of all the terms involving 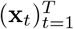, the outputs of the FIS suffice to compute the expectation of the objective function in Eq. (8), i.e., the inner E step, which results in:

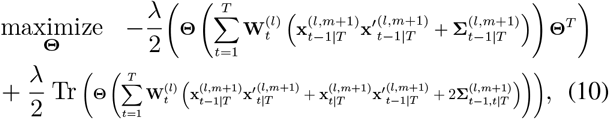

to obtain 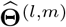. The solution has a closed-form given by:

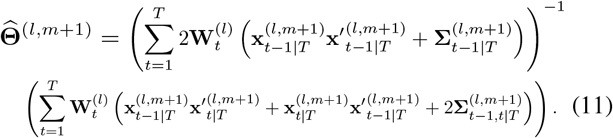

This process is repeated for *M* iterations for the inner EM and *L* iterations for the outer EM, until a convergence criterion is met. Algorithm 1 summarizes the FCSS algorithm.

**Algorithm 1.**
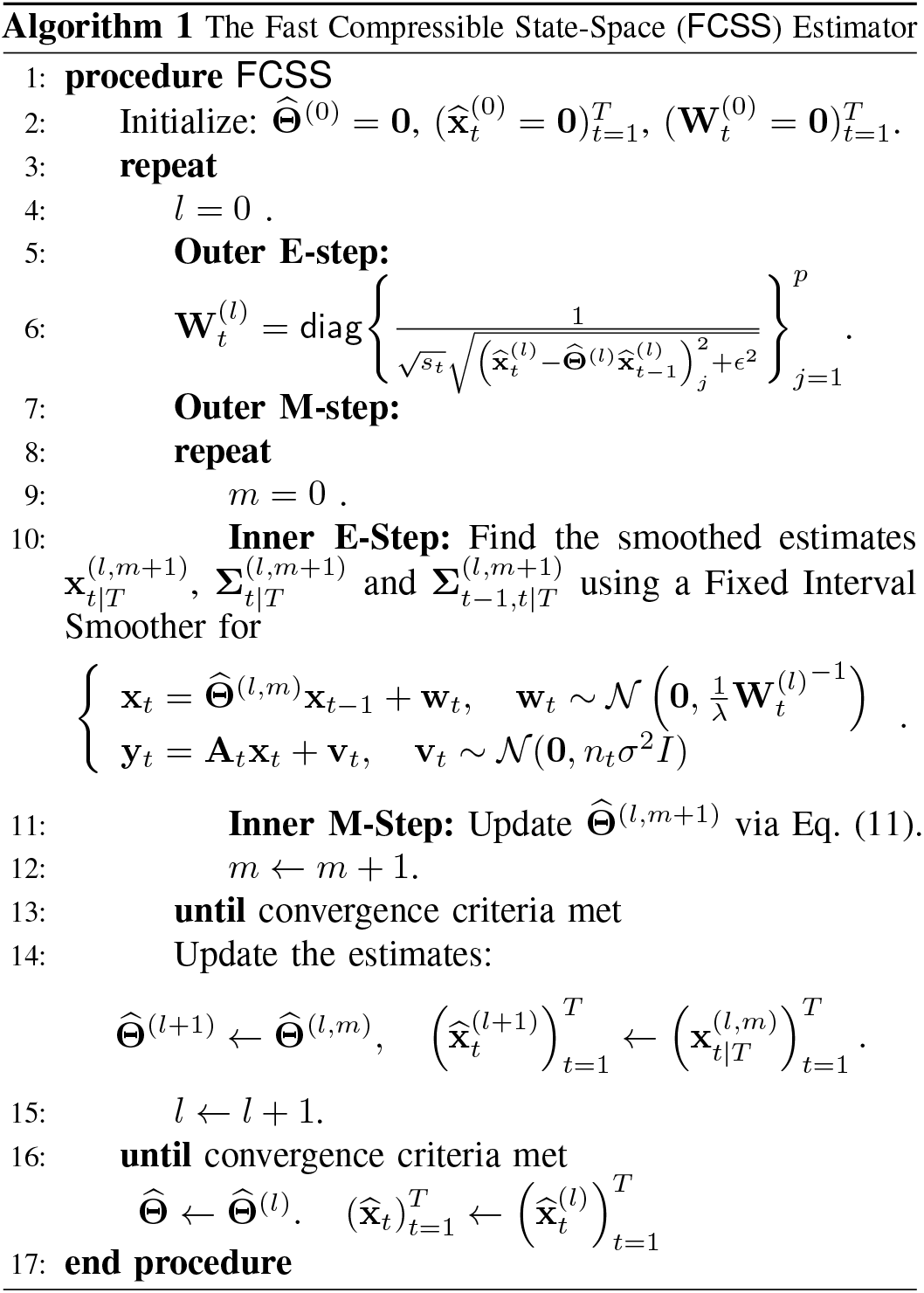
The Fast Compressible State-Space (FCSS) Estimator.

#### Remark 1.

By virtue of the FIS procedure, the compexity of the FCSS algorithm is *linear* in *T*, i.e., the observation duration. As we will show in Section III, this makes the FCSS algorithm scale favorably when applied to long data sets.

#### Remark 2.

In order to update**Θ** in the inner M-step given by E.(11), we have not specifically enforced the condition ∥**Θ**∥< 1 in the maximization step. This condition is required to obtain a convergent state transition matrix which results in the stability of the state dynamics. It is easy to verify that the set of matrices **Θ** satisfying ∥**Θ**∥< 1−*η*, is a closed convex set for small positive *η*;, and hence one can perform the maximization in (11) by projection onto this closed convex set. Alternatively, matrix optimization methods with operator norm constraints can be used [35]. We have avoided this technicality by first finding the global minimum and examining the largest eigenvalue. In the applications of interest in this paper which follow next, the largest eigenvalue has always been found to be less than 1.

### C. Experimental Procedures

*Surgery*: 2 hours before surgery, 0.1 cc dexamethasone (2 mg/ml, VetOne) was injected subcutaneously to reduce brain swelling during craniotomy. Anesthesia is induced with 4% isoflurane (Fluriso, VetOne) with a calibrated vaporizer (Matrx VIP 3000). During surgery, isoflurane level was reduced to and maintained at a level of 1.5%–2%. Body temperature of the animal is maintained at 36.0 degrees Celsius during surgery. Hair on top of head of the animal was removed using Hair Remover Face Cream (Nair), after which Betadine (Purdue Products) and 70% ethanol was applied sequentially 3 times to the surface of the skin before removing the skin. Soft tissues and muscles were removed to expose the skull. Then a custom designed 3D printed stainless headplate was mounted over left auditory cortex and secured with C&B-Metabond (Parkell). A craniotomy with a diameter of around 3.5 mm was then performed over left auditory cortex. A three layered cover slip was used as cranial window, which is made by gluing (NOA71, Norland Products) 2 pieces of 3 mm coverslips (64-0720 (CS-3R), Warner Instruments) with a 5 mm coverslip (64–0700 (CS-5R), Warner Instruments). Cranial window was quickly dabbed in kwik-sil (World Precision Instruments) before mounted 3 mm coverslips facing down onto the brain. After kwik-sil cured, Metabond was applied to secure the position of the cranial window. Synthetic Black Iron Oxide (Alpha Chemicals) was then applied to the hardened Metabond surface. 0.05 cc Cefazolin (1 gram/vial, West Ward Pharmaceuticals) was injected subcutaneously when entire procedure was finished. After the surgery the animal was kept warm under heat light for 30 minutes for recovery before returning to home cage. Medicated water (Sulfamethoxazole and Trimethoprim Oral Suspension, USP 200 mg/40 mg per 5 ml, Aurobindo pharms USA; 6 ml solution diluted in 100 ml water) substitute normal drinking water for 7 days before any imaging was performed.

*Awake two-photon imaging:* Spontaneous activity data of population of layer 2/3 auditory cortex (A1) neurons is collected from adult (3-month old) Thy1-GCaMP6s female mouse implanted with chronic window following the above procedure, using two-photon imaging. Acquisition is performed using a two-photon microscope (Thorlabs Bscope 2) equipped with a Vision 2 Ti:Sapphire laser (Coherent), equipped with a GaAsP photo detector module (Hamamatsu) and resonant scanners enabling faster high-resoluation scanning at 30–60 Hz per frame. The excitation wavelength was 920 nm. Regions (~ 300 μm^2^) within A1 were scanned at 30 Hz through a 20x, 0.95 NA water-immersion objective (Olympus). During imaging the animal was head-fixed and awake. The microscope was rotated 45 degrees and placed over the left A1 where window was placed. An average image of field of view was generated by choosing a time window where minimum movement of the brain was observed and used as reference image for motion correction using TurboReg plugin in ImageJ. GCaMP6s positive cells are selected manually by placing a ring like ROI over each identified cell. Neuropil masks were generated by placing a 20 *μ*m radius circular region over each cell yet excluding all cell soma regions. Traces of soma and neuropil were generated by averaging image intensity within respective masks at each time point. A ratio of 0.7 was used to correct for neuropil contamination.

*Cell-attached patch clamp recordings and two-photon imaging*: Recordings were performed in vitro in voltage clamp to simultaneously measure spiking activity and ∆*F*/*F*. Thalamocortical slices containing A1 were prepared as previously described [36]. The extracellular recording solution consisted of artificial cerebral spinal fluid (ACSF) containing: 130 NaCl, 3 KCl, 1.25 KH2PO4, 20 NaHCO3, 10 glucose, 1.3 MgSO4, 2.5 CaCl2 (pH 7.35-7.4, in 95% O2 5% CO2). Action potentials were recorded extracellularly in loose-seal cell-attached configuration (seal resistance typically 20–30 MΩ) in voltage clamp mode. Borosilicate glass patch pipettes were filled with normal ACSF diluted 10%, and had a tip resistance of ~3-5 MΩ in the bath. Data were acquired with a Multiclamp 700B patch clamp amplifier (Molecular Devices), low-pass filtered at 3-6 kHz, and digitized at 10 kHz using the MATLAB-based software. Action potentials were stimulated with a bipolar electrode placed in L1 or L23 to stimulate the apical dendrites of pyramidal cells (pulse duration 1-5 ms). Data were analyzed offline using MATLAB. Imaging was largely performed using a two-photon microscope (Ultima, Prairie Technologies) and a MaiTai DeepSee laser (SpectraPhysics), equipped with a GaAsP photo detector module (Hamamatsu) and resonant scanners enabling faster high-resoluation scanning at 30-60 Hz per frame. Excitation was set at 900 nm. Regions were scanned at 30 Hz through a 40x water-immersion objective (Olympus). Cells were manually selected as ring-like regions of interest (ROIs) that cover soma but exclude cell nuclei, and pixel intensity within each ROI was averaged to generate fluorescence over time and changes in fluorescence (∆*F*/*F*) were then calculated.

## III. Results

In this section, we study the performance of the FCSS estimator on simulated data as well real data from two-photon calcium imaging recordings of neuronal activity and sleep spindle detection from EEG.

### A. Application to Simulated Data

We first apply the FCSS algorithm to simulated data and compare its performance with the Basis Pursuit Denoising (BPDN) algorithm. The parameters are chosen as *p* = 200, *T* = 200, *s*_1_ = 8, *s*_2_ = 4, ∈ = 10^−10^, and **Θ** = 0.95**I**. We define the quantity 1 − *n*/*p* as the compression ratio. We refer to the case of *n_t_* = *p*, i.e., no compression, as the denoising setting. The measurement matrix **A** is an *n_t_* x *p* i.i.d. Gaussian random matrix, where *n_t_* is chosen such that 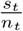 is a fixed ratio. An initial choice of 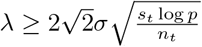 is made inspired by the theory of LASSO [37], which is further tuned using two-fold cross-validation.

Figures 1-(a) and 1-(b) show the estimated states as well as the innovations for different compression ratios for one sample component. In the denoising regime, all the innovations (including the two closely-spaced components) are exactly recovered. As we take fewer measurements, the performance of the algorithm degrades as expected. However, the overall structure of the innovation sequence is captured even for highly compressed measurements.

**Fig. 1.**
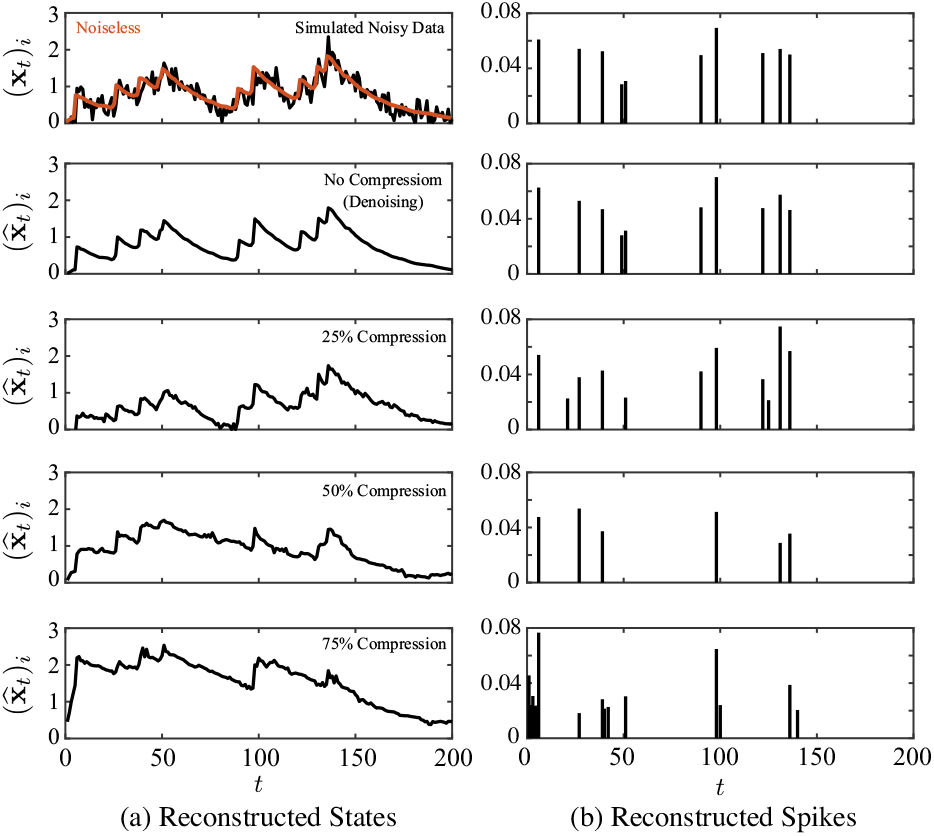
Reconstruction results of FCSS on simulated data vs. compression levels. (a) reconstructed states, (b) reconstructed spikes. The FCSS estimates degrade gracefully as the compression level increases.

Figure 2 shows the MSE comparison of the FCSS vs. BPDN, where the MSE is defined as 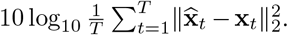 The FCSS algorithm significantly outperforms BPDN, especially at high SNR values. Figure 3 compares the performance of FCSS and BPDN on a sample component at a compression level of *n*/*p* = 2/3, in order to visualize the performance gain implied by Figure 3.

**Fig. 2.**
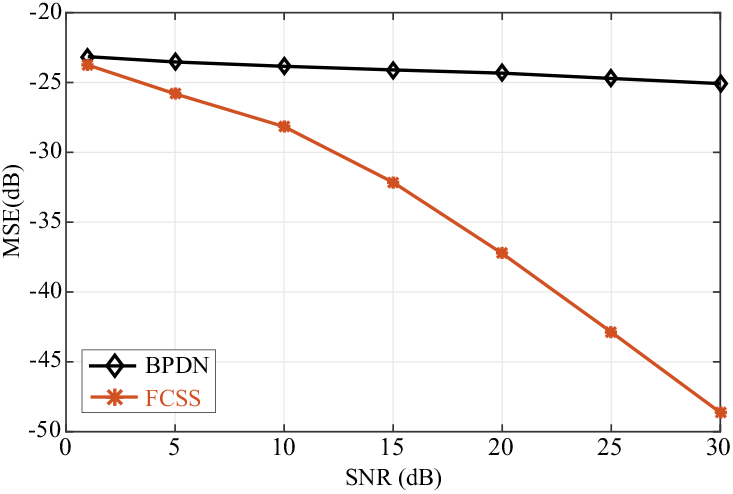
MSE vs. SNR comparison between FCSS and BPDN. The FCSS significantly outperforms the BPDN, even for moderate SNR levels.

**Fig. 3.**
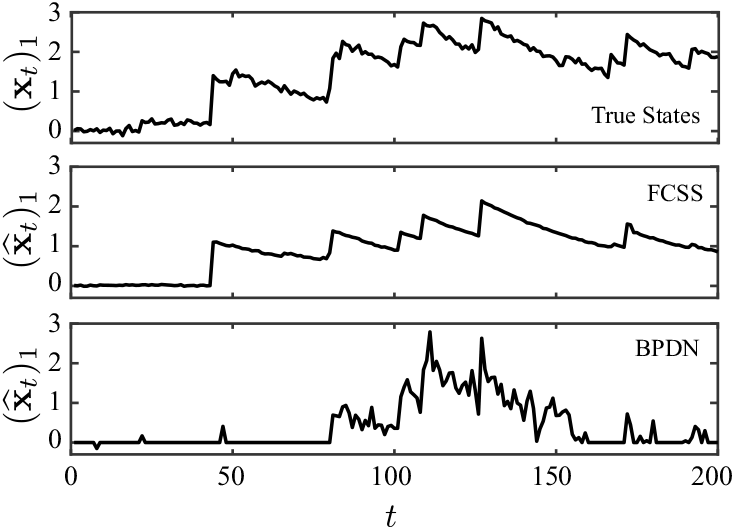
Example of state reconstruction results for FCSS and BPDN. Top: true states, Middle: FCSS state estimates, Bottom: BPDN estimates. The FCSS reconstruction closely follows the true state evolution, while the BPDN fails to capture the state dynamics.

Finally, Figure 4 shows the comparison of the estimated states for the entire simulated data in the denoising regime. As can be observed from the figure, the sparsity pattern of the states and innovations are captured while significantly denoising the observed states.

**Fig. 4.**
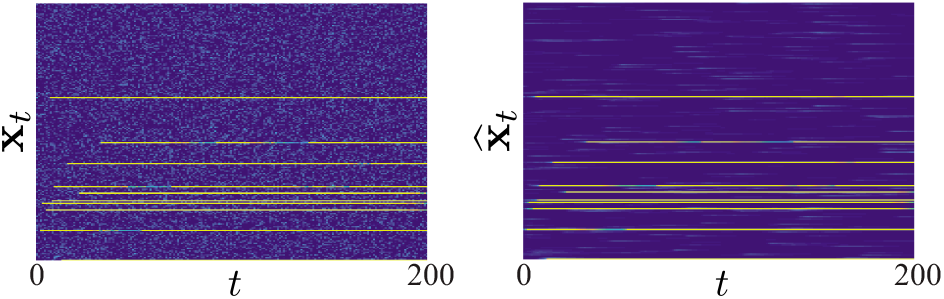
Raster plots of the observed and estimated states via FCSS. Left: noisy observations, Right: FCSS estimates. The FCSS significantly denoises the observed states.

### B. Application to Calcium Signal Deconvolution

Calcium imaging takes advantage of intracellular calcium flux to directly visualize calcium signaling in living neurons. This is done by using calcium indicators, which are fluorescent molecules that can respond to the binding of calcium ions by changing their fluorescence properties and using a fluorescence or two-photon microscope and a CCD camera to capture the visual patterns [38], [39]. Since spikes are believed to be the units of neuronal computation, inferring spiking activity from calcium recordings, referred to as calcium deconvolution, is an important problem in neural data analysis. Several approaches to calcium deconvolution have been proposed in the neuroscience literature, including model-free approaches such as sequential Monte Carlo methods [40] and model-based approaches such as non-negative deconvolution methods [5], [41]. These approaches require solving convex optimization problems, which do not scale well with the temporal dimension of the data. In addition, they lack theoretical performance guarantees and do not provide clear measures for assessing the statistical significance of the detected spikes.

In order to construct confidence bounds for our estimates, we employ recent results from high-dimensional statistics [42]. We first compute the confidence intervals around the outputs of the FCSS estimates using the node-wise regression procedure of [42], at a confidence level of 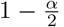. We perform the nodewise regression separately for each time *t*. For an estimate 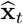, we obtain 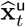 and 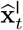 as the upper and lower confidence bounds, respectively. Next, we partition the estimates into small segments, starting with a local minimum (trough) and ending in a local maximum (peak). For the *i*^th^ component of the estimate, let *t*_min_ and *t*_max_ denote the time index corresponding to two such consecutive troughs and peaks. If the difference 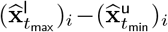 is positive, the detected innovation component is declared significant (i.e., spike) at a confidence level of 1 − α, otherwise it is discarded (i.e., no spike). We refer to this procedure as Pruned-FCSS (PFCSS).

We first apply the FCSS algorithm for calcium deconvolution in a scenario where the ground-truth spiking is recorded *in vitro* through simultaneous electrophysiology (cell-attached patch clamp) and two-photon calcium imaging. The calcium trace as well as the ground-truth spikes are shown for a sample neuron in Figure 5-(a). The FCSS denoised estimate of the states (black) and the detected spikes (blue) using 95% confidence intervals (orange hulls) and the corresponding quantities for the constrained f-oopsi algorithm [41] are shown in Figures 5-(b) and 5-(c), respectively. Both algorithms detect the large dynamic changes in the data, corresponding to the spikes, which can also be visually captured in this case. However, in doing so, the f-oopsi algorithm incurs a high rate of false positive errors, manifested as clustered spikes around the ground truth events. Similar to f-oopsi, most state-of-the-art model-based methods suffer from high false positive rate, which makes the inferred spike estimates unreliable. Thanks to the aforementioned pruning process based on the confidence bounds, the PFCSS is capable of rejecting the insignificant innovations, and hence achieve a lower false positive rate. One factor responsible for this performance gap can be attributed to the underestimation of the calcium decay rate in the transition matrix estimation step of f-oopsi. However, we believe the performance gain achieved by FCSS is mainly due to the explicit modeling of the sparse nature of the spiking activity by going beyond the Gaussian state-space modeling paradigm.

**Fig. 5.**
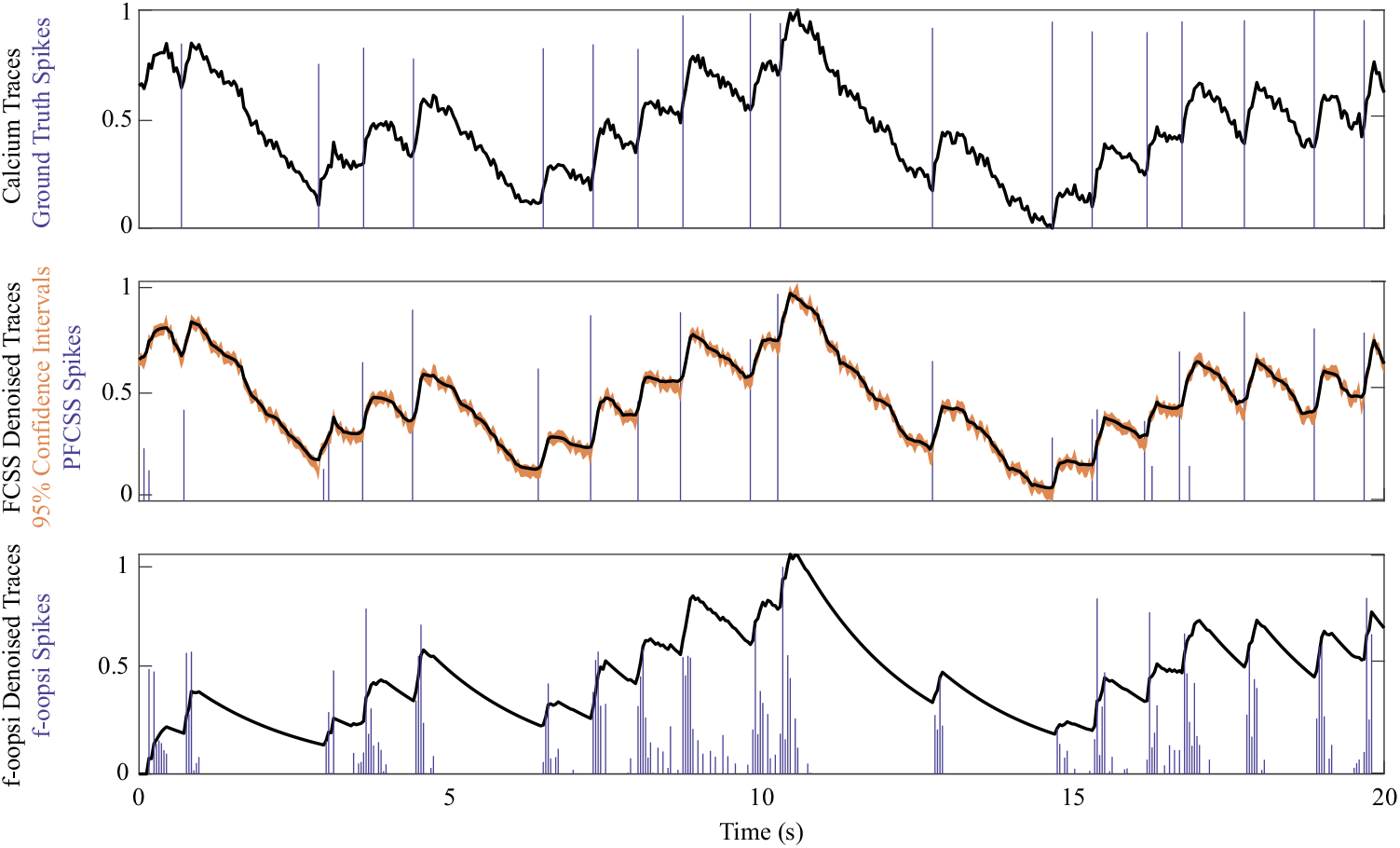
Ground-truth performance comparison between PFCSS and constrained f-oopsi. Top: the observed calcium traces (black) and ground-truth electrophysiology data (blue), Middle: PFCSS state estimates (black) with 95% confidence intervals (orange) and the detected spikes (blue), Bottom: the constrained f-oopsi state estimates (black) and the detected spikes (blue). The FCSS spike estimates closely match the ground-truth spikes with only a few false detections, while the constrained f-oopsi estimates contain significant clustered false detections.

Next, we apply the FCSS algorithm to large-scale *in vivo* calcium imaging recordings, for which the ground-truth is not available due to measurement constraints. The data used in our analysis was recorded from 219 spontaneously active neurons in mouse auditory cortex. The two-photon microscope operates at a rate of 30 frames per second. We chose *T* = 2000 samples corresponding to 1 minute for the analysis. We chose *p* = 108 well-separated neurons visually. We estimate the measurement noise variance by appropriate re-scaling of the power spectral density in the high frequency bands where the signal is absent. We chose a value of *∊* = 10^−10^. It is important to note that estimation of the measurement noise variance is critical, since it affects the width of the confidence intervals and hence the detected spikes. Moreover, we estimate the baseline fluorescence by averaging the signal over values within a factor of 3 standard deviations of the noise. By inspecting Eq.(4), one can see a trade-off between the choice of λ and the estimate of the observation noise variance *σ*^2^. We have done our analysis in both the compression regime, with a compression ratio of 1/3 (*n*/*p* = 2/3), and the denoising regime. The measurements in the compressive regime were obtained from applying i.i.d. Gaussian random matrices to the observed calcium traces. The latter is done to motivate the use of compressive imaging, as opposed to full sampling of the field of view.

Figure 6–(a) shows the observed traces for four selected neurons. The reconstructed states using FCSS in the compressive and denoising regimes are shown in Figures 6–(b) and –Figure 6(c), respectively. The 90% confidence bounds are shown as orange hulls. The FCSS state estimates are significantly denoised while preserving the calcium dynamics. Figure 7 shows the detected spikes using constrained f-oopsi and PFCSS in both the compressive and denoising regimes. Finally, Figure 8 shows the corresponding raster plots of the reconstructed spikes for the entire ensemble of neurons. Similar to the preceding application on ground-truth date, the f-oopsi algorithm detects clusters of spikes, whereas the PFCSS procedure results in sparser spike detection. This results in the detection of seemingly more active neurons in the raster plot. However, motivated by the foregoing ground-truth analysis, we believe that a large fraction of these detected spikes may be due to false positive errors. Strikingly, even with a compression ratio of 1/3 the performance of the PFCSS is similar to the denoising case. The latter observation corroborates the feasibility of compressed two-photon imaging, in which only a random fraction of the field of view is imaged, which in turn can result in higher acquisition rates.

**Fig. 6.**
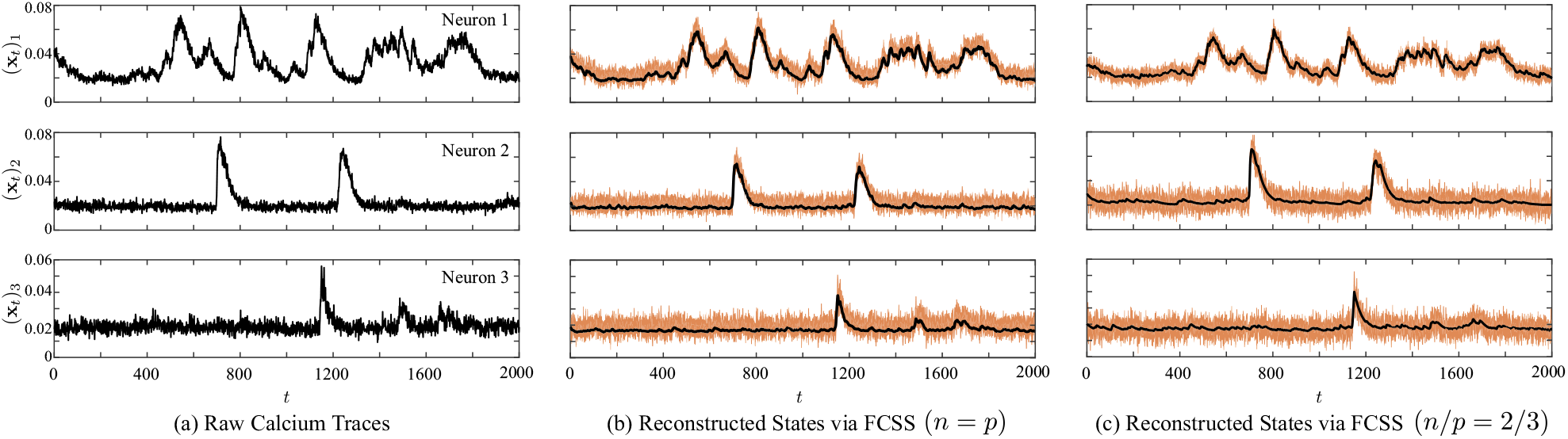
Performance of FCSS on large-scale calcium imaging data. Top, middle and bottom rows correspond to three selected neurons labeled as Neuron 1, 2, and 3, respectively. (a) raw calcium traces, (b) FCSS reconstruction with no compression, (c) FCSS reconstruction with 2/3 compression ratio. Orange hulls show 95% confidence intervals. The FCSS significantly denoises the observed traces in both the uncompressed and compressed settings.

**Fig. 7.**
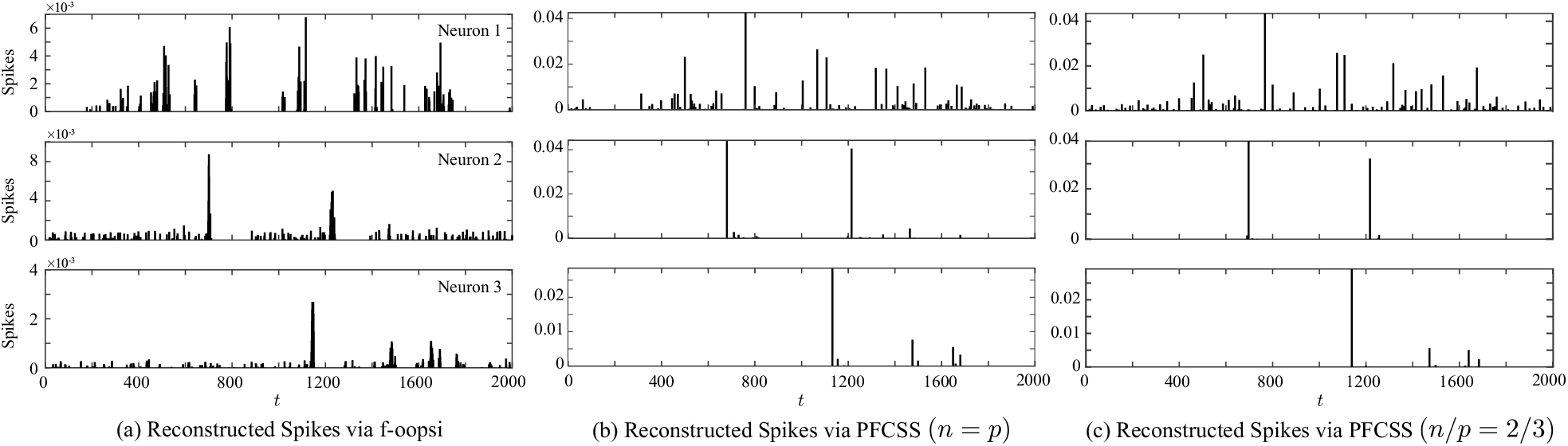
Reconstructed spikes of PFCSS and constrained f-oopsi from large-scale calcium imaging data. Top, middle and bottom rows correspond to three selected neurons labeled as Neuron 1, 2, and 3, respectively. (a) constrained f-oopsi spike estimates, (b) PFCSS spike estimates with no compression, (c) PFCSS spike estimates with 2/3 compression ratio. The PFCSS estimates in both the uncompressed and compressed settings are sparse in time, whereas the constrained f-oopsi estimates are in the form of clustered spikes.

**Fig. 8.**
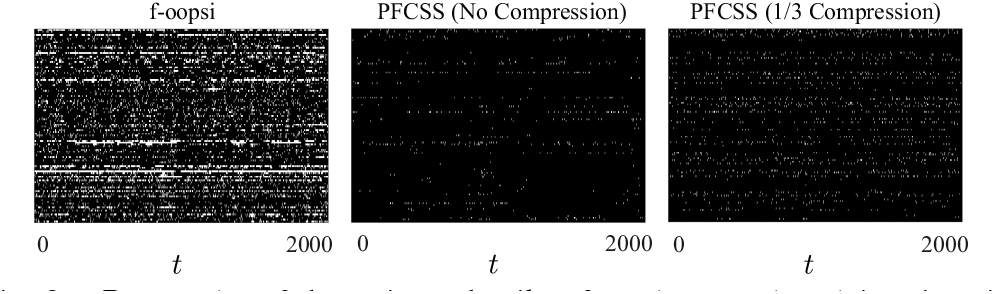
Raster plot of the estimated spikes from large-scale calcium imaging data. Left: the constrained f-oopsi estimates, Middle: PFCSS estimates with no compression, Right: PFCSS estimates with a ⅓ compression ratio. The PFCSS estimates are spatiotemporally sparse, whereas the constrained f-oopsi outputs temporally clustered spike estimates.

In addition to the foregoing discussion on the comparisons in Figure 5, 6, 7, and 8, two remarks are in order. First, the iterative solution at the core of FCSS is linear in the observation length and hence significantly faster than the batch-mode optimization procedure used for constrained f-oopsi. Our comparisons suggest that the FCSS reconstruction is at least 3 times faster than f-oopsi for moderate data sizes of the order of tens of minutes. Moreover, the vector formulation of FCSS allows for easy parallelization (e.g., via GPU implementations), which allows simultaneous processing of ROI’s without losing speed. Second, using only about two-thirds of the measurements achieves similar results by FCSS as using the full measurements.

### C. Application to Sleep Spindle Detection

In this section we use compressible state-space models in order to model and detect sleep spindles. A sleep spindle is a burst of oscillatory brain activity manifested in the EEG that occurs during stage 2 non-rapid eye movement (NREM) sleep. It consists of stereotypical 12–14 Hz wave packets that last for at least 0.5 seconds [43]. The spindles occur with a rate of 2–5% in time, which makes their generation an appropriate candidate for compressible dynamics. Therefore, we hypothesize that the spindles can be modeled using a combination of few echoes of the response of a second order compressible state-space model. As a result, the spindles can be decomposed as sums of modulated sine waves.

In order to model the oscillatory nature of the spindles, we consider a second order autoregressive (AR) model where the pole locations are given by 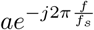 and 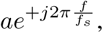 where 0 < *a* < 1 is a positive constant controlling the effective duration of the impulse response, *f_s_* is the sampling frequency and *f* is a putative frequency accounting for the dominant spindle frequency. The equivalent state-space model for which the MAP estimation admits the FCSS solution is therefore:

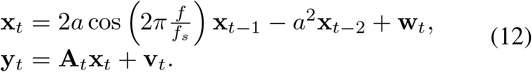

Note that impulse response of the state-space dynamics is given by 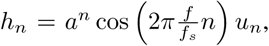, which is a stereotypical decaying sinusoid. By defining the augmented state 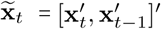, Eq.(12) can be expressed in the canonical form:

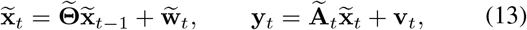

where 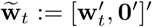,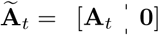 and

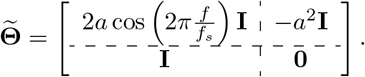

Eq. (10) can be used to update 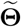 in the M step of the FCSS algorithm. However, 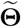 has a specific structure in this case, determined by *a* and *f*, which needs to be taken into account in the optimization step. Let 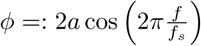 and 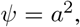, and let

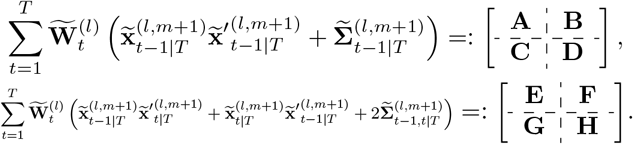

Then, Eq. (10) is equivalent to maximizing

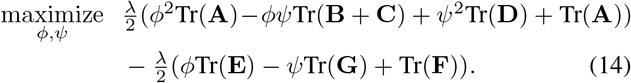

subject to 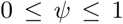 and 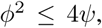, which can be solved using interior point methods. In our implementation, we have imposed additional priors of *f* ~ Uniform(12,14) Hz and a ~ Uniform(0.95,0.99), which simplifies the constraints on *ϕ* and *ψ* to

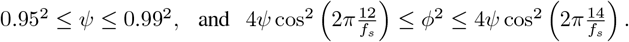

Given the convexity of the cost function in (14), one can conclude from the KKT conditions that if the global minimum is not achieved inside the region of interest it must be achieved on the boundaries.

Figure 9, top panel, shows two instances of simulated spindles (black traces) with parameters *f_s_* = 200 Hz, *f* = 13 Hz and *a* = 0.95, with the ground truth events generating the wave packets shown in red. The middle panel shows the noisy version of the data with an SNR of –7.5 dB. The noise was chosen as white Gaussian noise plus slowly varying (2 Hz) oscillations to resemble the slow oscillations in real EEG data. As can be observed the simulated signal exhibits visual resemblance to real spindles, which verifies the hypothesis that spindles can be decomposed into few combinations of wave packets generated by a second order AR model. The third panel, shows the denoised data using FCSS, which not only is successful in detecting the the ground-truth dynamics (red bars), but also significantly denoises the data.

**Fig. 9.**
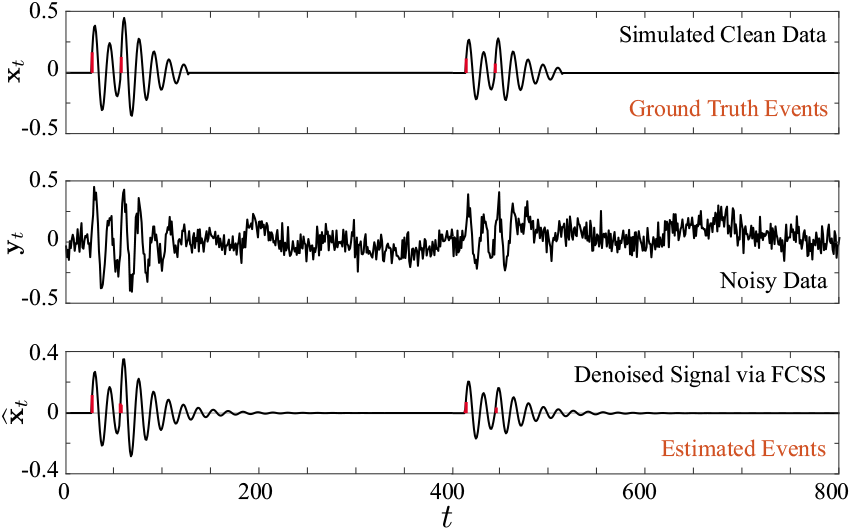
Performance of FCSS on simulated spindles. Top: simulated clean data (black) and ground-truth spindle events (red), Middle: simulated noisy data, Bottom: the denoised signal (black) and deconvolved spindle events (red). The FCSS estimates are significantly denoised and closely match the ground-truth data shown in the top panel.

We next apply FCSS to real EEG recordings from stage-2 NREM sleep. Manual scoring of sleep spindles can be very time-consuming, and achieving accurate manual scoring on a long-term recording is a highly demanding task with the associated risk of decreased diagnosis. Although automatic spindle detection would be attractive, most available algorithms sensitive to variations in spindle amplitude and frequency that occur between both subjects and derivations, reducing their effectiveness [44], [45]. Moreover most of these algorithms require significant pre- and post-processing and manual tuning. Examples include algorithms based on Empirical Mode Decomposition (EMD) [46], [47], [48], data-driven Bayesian methods [49], and machine learning approaches [50], [51]. Unlike our approach, none of the existing methods consider modeling the generative dynamics of spindles, as transient sparse events in time, in the detection procedure.

The data used in our analysis is part of the recordings in the DREAMS project [52], recorded using a 32-channel polysomnograpgh. We have used the EEG channels in our analysis. The data was recorded at a rate of *f_s_* = 200 Hz for 30 minutes. The data was scored for sleep spindles independently by two experts. We have used expert annotations to separate regions which include spindles for visualization purposes. For comparison purposes, we use a bandpass filtered version of the data within 12–14 Hz, which is the basis of several spindle detection algorithms [45], [53], [54], hallmarked by the widelyused bandpass filtered root-mean-square (RMS) method [45].

Figure 10 shows the detection results along with the bandpass filtered version of the data for two of the EEG channels. The red bars show the expert markings of the onset and offset of the spindle events. The FCSS simultaneously captures the spindle events and suppressed the activity elsewhere, whereas the bandpass filtered data produces significant activity in the 12–14 Hz throughout the observation window, resulting in high false positives. To quantify this observation, we have computed the ROC curves of the FCSS and bandpass filtering followed by root mean square (RMS) computation in Figure 11, which confirms the superior performance of the FCSS algorithm over the data set. The annotations of one of the experts has been used for as the ground truth benchmark.

**Fig. 10.**
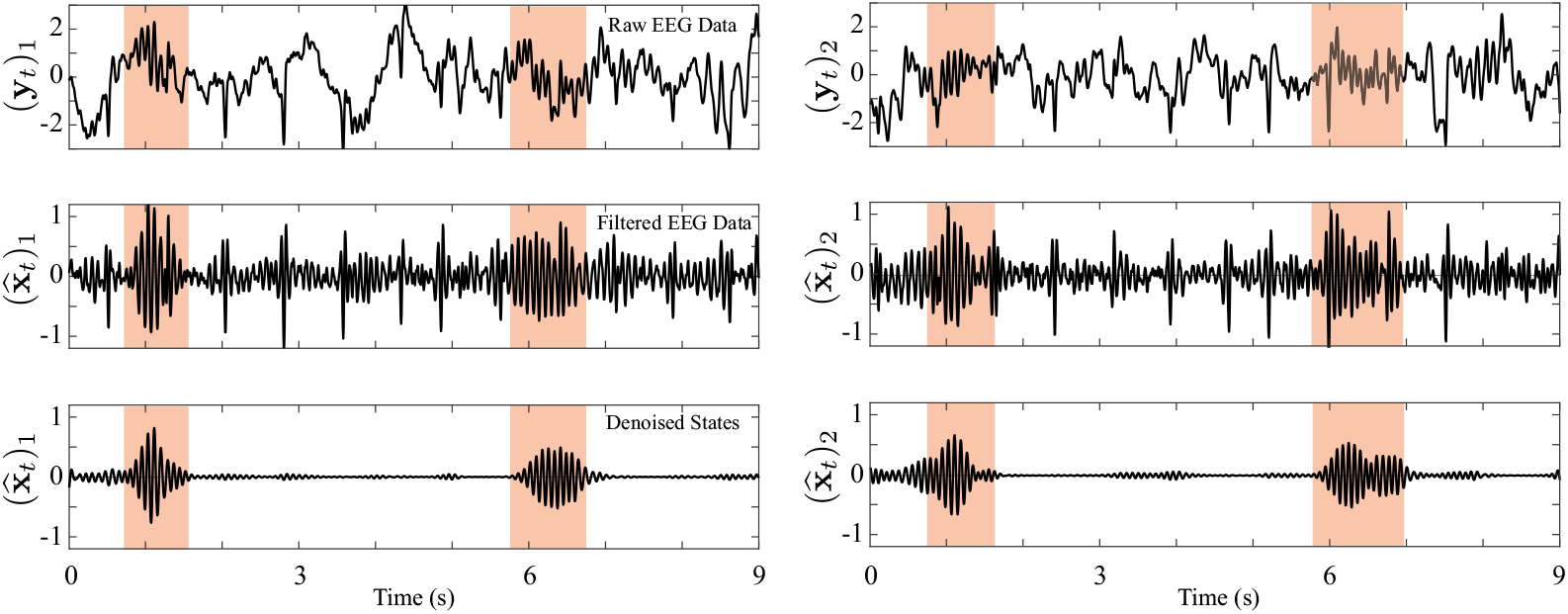
Performance comparison between FCSS and band-pass filtered EEG data. Left and right panels correspond to two selected electrodes labeled as 1 and 2, respectively. The orange blocks show the extent of the detected spindles by the expert. Top: raw EEG data, Middle: band-pass filtered EEG data in the 12–14 Hz band, Bottom: FCSS spindle estimates. The FCSS estimates closely match the expert annotations, while the band-pass filtered data contains significant signal components outside of the orange blocks.

**Fig. 11.**
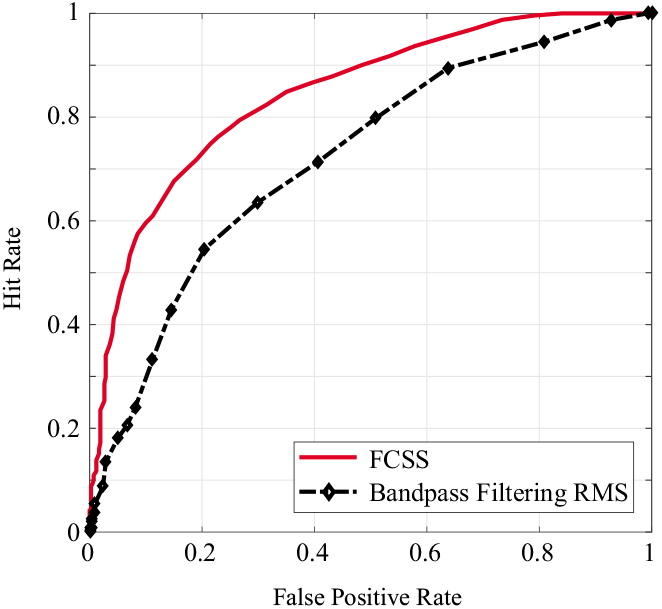
The ROC curves for FCSS (solid red) and bandpass-filtered RMS (dashed black). The FCSS outperforms the widely-used band-pass filtered RMS method as indicated by the ROC curves.

## IV. Discussion

In this section, we discuss the implication of our techniques in regard to the application domains as well as existing methods.

### A. Connection to existing literature in sparse estimation

Contrary to the traditional compressive sensing, our linear measurement operator does not satisfy the RIP [21], despite the fact that A_*t*_’s satisfy the RIP. Nevertheless, we have extended the near-optimal recovery guarantees of CS to our compressible state-space estimation problem via Theorem 1. Closely related problems to our setup are the super-resolution and sparse spike deconvolution problems [55], [56], in which abrupt changes with minimum separation in time are resolved in fine scales using coarse (lowpass filtered) frequency information, which is akin to working in the compressive regime.

Theoretical guarantees of CS require the number of measurements to be roughly proportional to the sparsity level for stable recovery [37]. These results do not readily generalize to the cases where the sparsity lies in the dynamics, not the states per se. Most of the dynamic compressive sensing techniques such as Kalman filtered compressed sensing, assume partial information about the support or estimate them in a greedy and often ad-hoc fashion [16], [17], [18], [19], [20]. As another example, the problem of recovering discrete signals which are approximately sparse in their gradients using compressive measurements, has been studied in the literature using Total Variation (TV) minimization techniques [57], [31]. For one-dimensional signals, since the gradient operator is not orthonormal, the Frobenius operator norm of its inverse grows linearly with the discretization level [57]. Therefore, stability results of TV minimization scale poorly with respect to discretization level. In higher dimensions, however, the fast decay of Haar coefficients allow for near-optimal theoretical guarantees [58]. A major difference of our setup with those of CS for TV-minimization is the *structured* and *causal* measurements, which unlike the non-causal measurements in [57], do not result in an overall measurement matrix satisfying RIP. We have considered dynamics with convergent transition matrices, in order to generalize the TV minimization approach. To this end, we showed that using the state-space dynamics one can infer temporally global information from local and causal measurements. Another closely related problem is the fused lasso [59] in which sparsity is promoted both on the covariates and their differences.

### B. Application to calcium deconvolution

In addition to scalability and the ability to detect abrupt transitions in the states governed by discrete events in time (i.e., spikes), our method provides several other benefits compared to other spike deconvolution methods based on state-space models, such as the constrained f-oopsi algorithm. First, our sampling-complexity trade-offs are known to be optimal from the theory of compressive sensing, whereas no performance guarantee exists for constrained f-oopsi. Second, we are able to construct precise confidence intervals on the estimated states, whereas constrained f-oopsi does not produce confidence intervals over the detected spikes. A direct consequence of these confidence intervals is estimation of spikes with high fidelity and low false alarm. Third, our comparisons suggest that the FCSS reconstruction is at least 3 times faster than f-oopsi for moderate data sizes of the order of tens of minutes. Finally, our results corroborate the possibility of using compressive measurement for reconstruction and denoising of calcium traces. From a practical point of view, a compressive calcium imaging setup can lead to higher scanning rate as well as better reconstructions, which allows monitoring of larger neuronal populations [60]. Due to the structured nature of our sampling and reconstruction schemes, we can avoid prohibitive storage problems and benefit from parallel implementations.

### C. Application to sleep spindle detection

Another novel application of our modeling and estimating framework is to case sleep spindle generation as a second-order dynamical system governed by compressive innovations, for which FCSS can be efficiently used to denoise and detect the spindle events. Our modeling framework suggest that spectrotemporal spindle dynamics cannot be fully captured by just pure sinusoids via bandpass filtering, as the data consistently contains significant 12–14 Hz oscillations almost everywhere (See Figure 10). Therefore, using the bandpass filtered data for further analysis purposes clearly degrades the performance of the resulting spindle detection and scoring algorithms. The FCSS provides a robust alternative to bandpass filtering in the form of model-based denoising.

In contrast to state-of-the-art methods for spindle detection, our spindle detection procedure requires minimal pre- and post-processing steps. We expect similar properties for higher order AR dynamics, which form a useful generalization of our methods for deconvolution of other transient neural signals. In particular, K-complexes during the stage 2 NREM sleep form another class of transient signals with high variability. A potential generalization of our method using higher order models can be developed for simultaneous detection of K-complexes and spindles.

## V. Conclusion

In this paper, we considered estimation of compressible state-space models, where the state innovations consist of compressible discrete events. For dynamics with convergent state transition dynamics, using theory of compressed sensing we provided an optimal error bound and stability guarantees for the dynamic 𝓁_1_-regularization algorithm which is akin to the MAP estimator for a Laplace state-space model. We also developed a fast and low-complexity iterative algorithm, namely FCSS, for estimation of the states as well as their transition matrix. We further verified the validity of our theoretical results through simulation studies as well as application to spike deconvolution from calcium traces and detection of sleep spindles from EEG data. Our methodology has two unique major advantages: first, we have proven theoretically why our algorithm performs well, and characterized its error performance. Second, we have developed a fast algorithm, with guaranteed convergence to a solution of the deconvolution problem, which for instance, is ~ 3 times faster than the widely-used f-oopsi algorithm in calcium deconvolution applications.

While we focused on two specific application domains, our modeling and estimation techniques can be generalized to apply to broader classes of signal deconvolution problems: we have provided a framework to model transient phenomena which are driven by sparse generators in time domain, and whose event onsets are of importance. Examples include heart beat dynamics and rapid changes in the covariance structure of neural data (e.g., epileptic seizures). In the spirit of easing reproducibility, we have made the MATLAB implementation of our algorithm publicly available [28].

## VI Acknowledgments

This work was supported in part by the National Science Foundation Award No. 1552946 and the National Institutes of Health Awards No. R01-DC009607 and U01-NS090569.

## VII Appendices

### A. Proof of Theorem 1

*Proof*: The main idea behind the proof is establishing appropriate cone and tube constraints [30]. In order to avoid unnecessary complications we assume *s*_1_ ≫ *s*_2_ =…= *s_T_* and *n*_1_ ≫*n*_2_=… =*n_T_*. (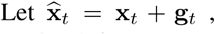 *t* ∈ [*T*] be an arbitrary solution to the primal form (3). We define z_*t*_ = x_*t*_ − *θ*x_*t*−1_ and (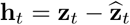 for *t* ∈ [*T*].For a positive integer *p*, let [*p*]: = {1, 2,…,*p*}. For an arbitrary set *V* ⊂ [*p*], x_*v*_ denotes the vector x restricted to the indices in *V*, i.e. all the components outside of *V* set to zero. We can decompose (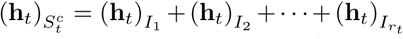, where *r_t_* = ⌊*p*/4*s_t_*⌋, and (h_*t*_)_*I*_1__ is the 4*s_t_*-sparse vector corresponding to 4*s_t_* largest-magnitude entries remaining in (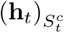, (h_*t*_)_*I*_2__ is the 4*s_t_*-sparse vector corresponding to 4*s* largest-magnitude entries remaining in (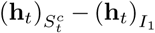 and so on. By the optimality of (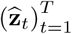 we have

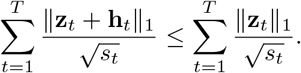

Using several instances of triangle inequality we have

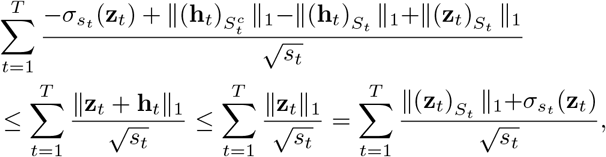

which after re-arrangement yields the cone condition given by

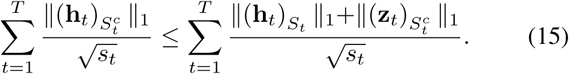

Also, by the definition of partitions 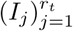 we have

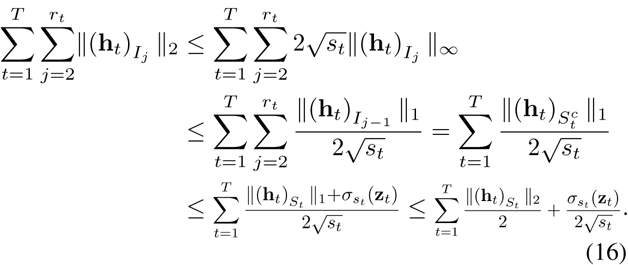

Moreover, using the feasibility of both x_*t*_ and 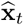 we have the tube constraints

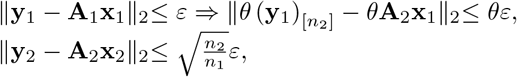

from which we conclude ∥y_2_ − θ (y_1_)_[*n*_2_]_ − A_2_z_2_∥_2_ ≤ (1 + *θ*)*ε*. Similarly 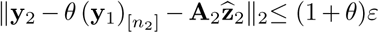. Therefore the triangle inequality yields ∥A_2_h_2_∥_2_≤ 2(1 + *θ*)*ε*. Similarly for all *t* ∈ [*T*]\{2}, we have the tighter bound

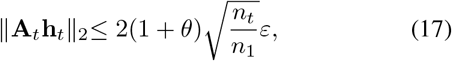

which is a consequence of having fewer measurements for *t* ∈ [*T*]\{2}. In conjunction, (15), (16), and (17) yield

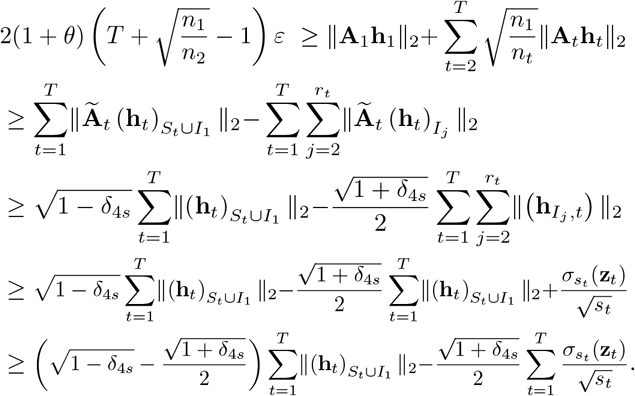

Therefore after rearrangement for *δ*_4*s*_ < 1/3

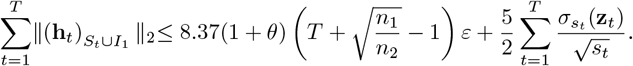

Next, using (16) yields

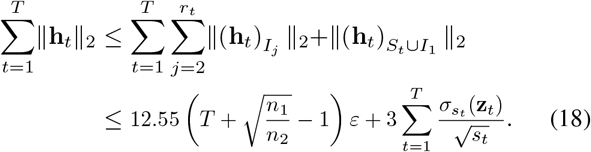

By definition we have h_*t*_ = g_*t*_ − *θ*g_*t*-1_ for *t* ∈ [*T*] with g_0_ = 0. Therefore by induction we have 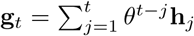 or in matrix form

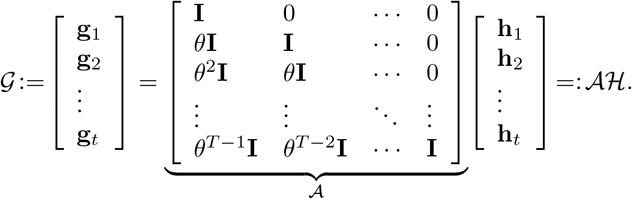

Using several instances of the triangle inequality we get:

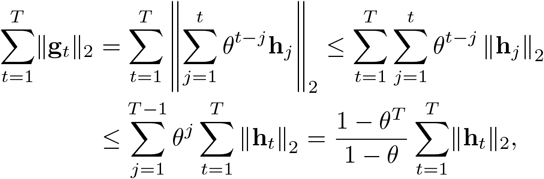

which in conjunction with (18) completes the proof.

### B. The Expectation Maximization Algorithm

In this section we give a short overview of the EM algorithm and its connection to iteratively re-weighted least squares (IRLS) algorithms. More details can be found in [33] and the references therein. Given the observations y, the goal of the EM algorithm is to find the ML estimates of a set of parameters **Θ** by maximizing the likelihood 𝔏(**Θ**): = *p*(y|**Θ**). Such maximization problems are typically intractable, but often become significantly simpler by introducing a latent variable u. The EM algorithm connects solving the ML problem to maximizing 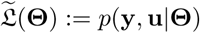, if one knew u.

Consider the state-space model:

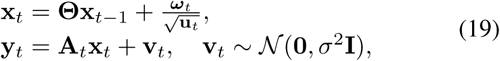

where 𝜔 ~ 𝒩(0, **I**), u_t_ is a positive i.i.d. random vector, and the square root operator and division of the two vectors are understood as element-wise operations. Let 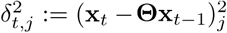 for j = 1,2,…,*p*. For an appropriate choice of the density of (u_*t*_)_*j*_ denoted by *p_u_*(.), we have [33]:

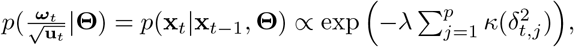

where

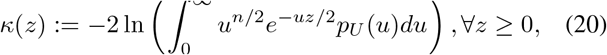

and k′(*z*) is a completely monotone function [61]. Random vectors of the form 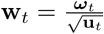 are known as Normal/Independent [61]. Note that a choice of 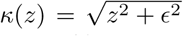 results in the *∊*-perturbed Laplace distributions used in our model [33]. Given *T* observations 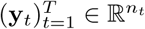 and conditionally independent samples 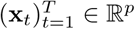, we denote the objective function of the MAP estimator by 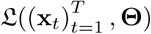, that is 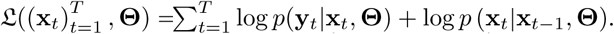 Consider the current estimates 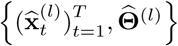 at iteration *l*. Then:

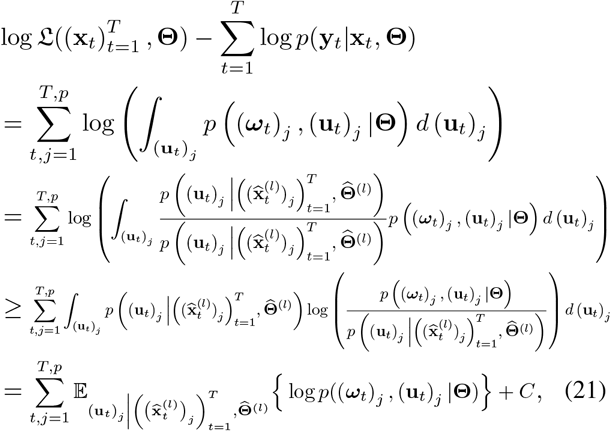

where the inequality follows from Jensen’s inequality and the constant *C* accounts for terms which do not depend on **Θ**. The so called Q-function is defined as:

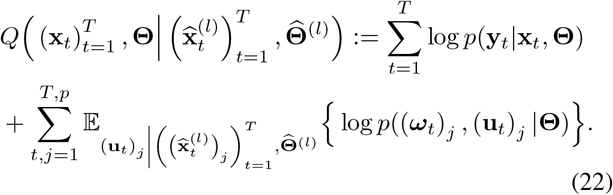

The EM algorithm maximizes the lower bound given by the Q-function of (22) instead of the log-likelihood itself. Moreover for all *t* ∈ [*T*], *j* ∈ [*p*] and 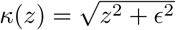 we have [61]:

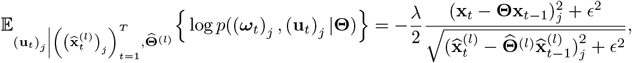

which after replacement results in the state-space model given by (9). This expectation gets updated in the outer EM loop using the *final* outputs of the inner loop. The outer EM algorithm can thus be summarized as forming the Q-function (E-step) and maximizing over **Θ** (M-step), which is known to converge to a stationary point due to its ascent property [33]. As discussed in Section II-B, the outer M-step is implemented by another instance of the EM algorithm by alternating between Fixed Interval Smoothing (E-step) and updating **Θ** (M-step).

